# INTEGRATION OF ENGINEERED “SPARK-CELL” SPHEROIDS FOR OPTICAL PACING OF CARDIAC TISSUE

**DOI:** 10.1101/2021.01.25.428177

**Authors:** Christianne Chua, Julie Han, Weizhen Li, Wei Liu, Emilia Entcheva

## Abstract

Optogenetic methods for pacing of cardiac tissue can be realized by direct genetic modification of the cardiomyocytes to express light-sensitive actuators, such as channelrhodopsin-2, ChR2, or by introduction of light-sensitized non-myocytes that couple to the cardiac cells and yield responsiveness to optical pacing. In this study, we engineer three-dimensional “spark cells” spheroids, composed of ChR2-expressing human embryonic kidney cells, and characterize their morphology as function of cell density and time. These “spark-cell” spheroids are then deployed to demonstrate site-specific optical pacing of human stem-cell-derived cardiomyocytes (hiPSC-CMs) in 96-well format using non-localized light application and all-optical electrophysiology. We show that the spheroids can be handled using liquid pipetting and can confer optical responsiveness of cardiac tissue earlier than direct viral or liposomal genetic modification of the cardiomyocytes, with 24% providing reliable stimulation of the iPSC-CMs within 6 hours and >80% within 24 hours. Our results demonstrate a scalable, cost-effective method to achieve contactless optical stimulation of cardiac cell constructs that can be integrated in a robotics-amenable workflow for high-throughput drug testing.

**GRAPHICAL ABSTRACT:** 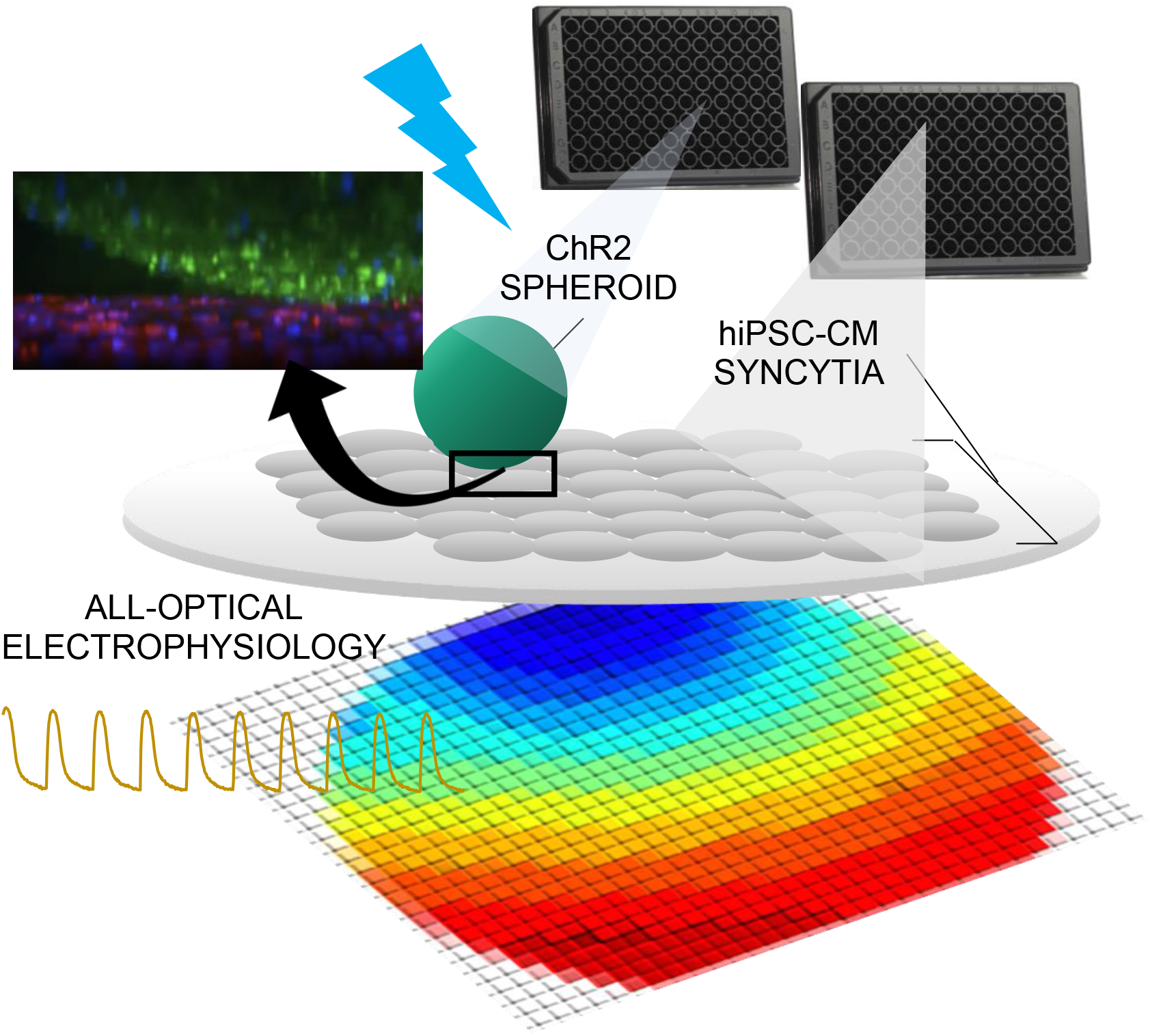

## Introduction

Traditionally, cardiac tissue is stimulated using electrodes delivering electrical pulses. Such electrodes require physical contact and cannot easily be deployed for multisite stimulation. Advances in optogenetics [1; 2; 3; 4] present an alternative - pacing tissue using light. Optogenetic modification via infection or transfection to introduce light-sensitive ion channels or opsins, such as channelrhodopsin-2 (ChR2), in cardiomyocytes allows pacing by light that offers certain benefits over electrical stimulation [5]. Optogenetic rhythm control has been deployed at the whole heart in a variety of studies[6; 7; 8; 9; 10; 11]. When combined with human induced pluripotent stem-cell-derived cardiomyocytes, hiPSC-CMs, in vitro, the contactless and scalable light-based optogenetic approaches hold promise to aid high-throughput (HT) capabilities for functional drug cardiotoxicity testing or other aspects of drug development [12; 13; 14; 15]. All-optical electrophysiology is poised to accelerate and streamline drug development [16; 17; 18; 19].

Direct transduction methods generally require several days for the cardiomyocytes to express the opsin and to become responsive to the pacing rhythm of blue-light pulses [14; 15; 20; 21; 22]. When done on site, they require special institutional protocols for handling recombinant DNA. On the other hand, it has been demonstrated that dedicated ChR2-expressing non-myocytes can be used as a driver to pace nontransduced myocytes by the so called “tandem-cell-unit” (TCU) approach[18; 21; 23], **Figure 1**. These studies highlighted a benefit of reducing the energy needed for pacing by spatially aggregating the light-responsive cells by cell patterning [21]. This approach can simplify the requirements for using optogenetic pacing (no genetic modification will be required on site) and may shorten the time needed to achieve light responsiveness for HT drug testing applications.

**FIGURE 1:**
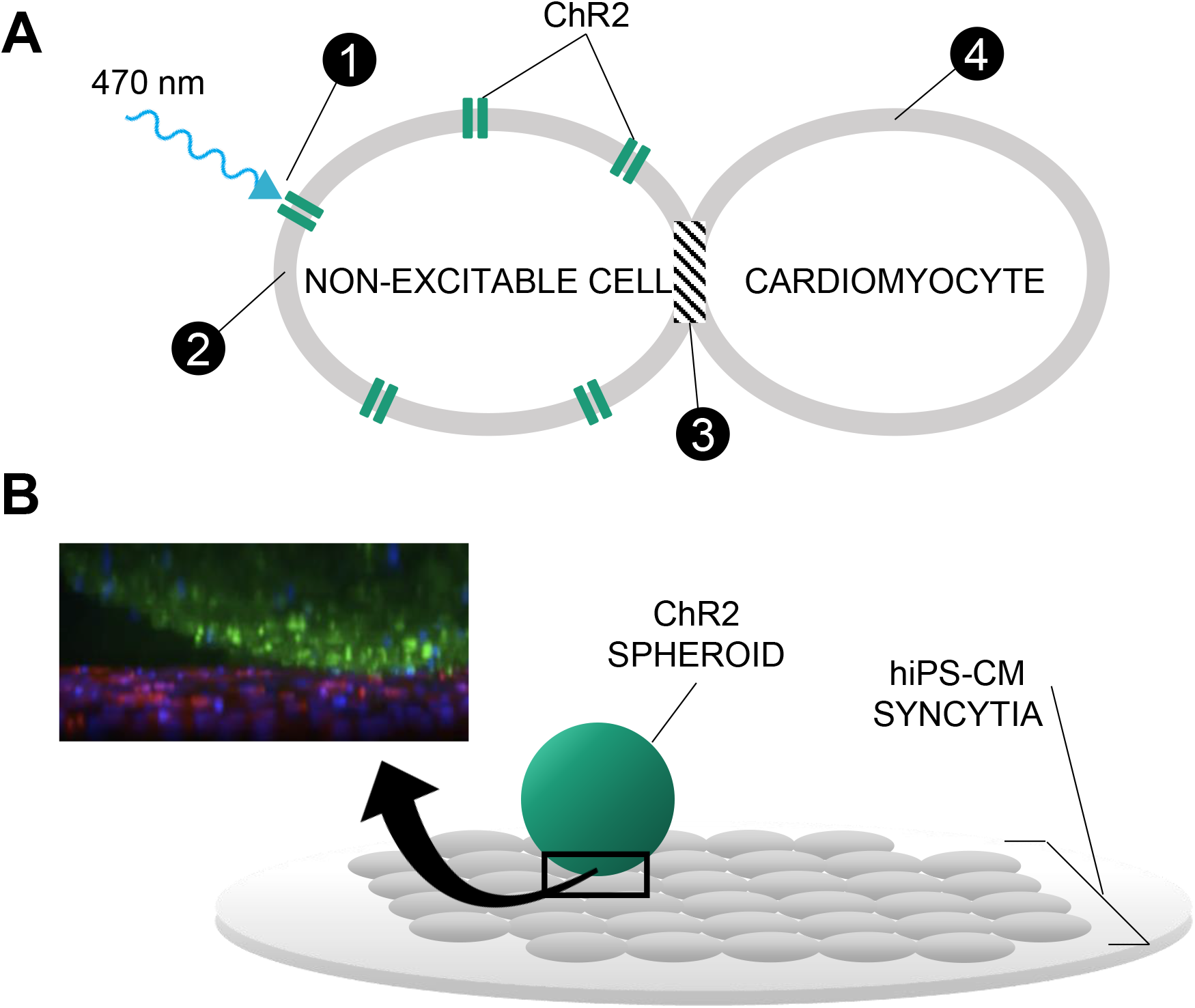
TANDEM-CELL-UNIT APPROACH USING 3D SPARK CELL SPHEROIDS. A: Illustrated is the tandem-cell-unit (TCU) approach. (1) Blue light (470 nm) induces conformational change in ChR2 expressed in a non-excitable cell resulting in (2) inward current and an elevated membrane potential in the non-excitable cell. Because there exists (3) some form of coupling between the non-excitable and cardiac cells, (4) the system evokes an action potential and subsequent contraction in the cardiomyocytes. B: Shown is the deployment of TCU using a spheroid structure and cardiac tissue – a schematic representation of a ChR2-expressing three-dimensional cell spheroid on top of a two-dimensional layer of hiPS-CM tissue. Confocal microscopy captured the relative positioning (ChR2-HEKs labeled green and hiPSC-CMs labeled red) as shown in the inset.

In this study, we demonstrate the manufacturing of three-dimensional constructs (spheroids) of ChR2-expressing “spark-cells” and characterize their integration and ability to stimulate syncytia of hiPS-CMs by the TCU strategy. The motivation for this study was two-fold: 1) develop a modular system for contactless pacing that is scalable and amenable to robotic handling; and 2) avoid any genetic manipulation of the cardiomyocytes and demonstrate faster timecourse of “spark-cell” spheroid-mediated conferment of optical pacing compared to direct transduction techniques. Our intent for these “spark-cell” spheroids is to be ultimately used as a “reagent”, fitting into a manufacturing workflow with robotic handling, long-term storage, transportation, and reliable deployment in HT drug testing applications.

## Materials and Methods

### ChR2-HEK SPHEROID ASSEMBLY

Previously, we have developed an immortal HEK cell line expressing ChR2 with a YFP tag [23] using Addgene construct pcDNA3.1 /hChR2(H 134R)-EYFP generously deposited of Karl Deisseroth. ChR2-HEKs were thawed, plated and expanded in standard T-75 flasks in a Dulbecco’s Modified Eagle Medium (DMEM) supplied with 10% fetal bovine serum (FBS) and 1% penicillin-streptomycin. ChR2-HEKs were lifted from the T-75 tissue culture flasks by 0.05% trypsin in Hanks’ Balanced Salt Solution (HBSS) following phosphate-buffered solution (PBS) rinse. After the resulting suspension underwent centrifugation, it was resuspended in a volume of DMEM appropriate to yield spheroids on the order of 10^4^ cells per well of a 96-well microplate. Special ultra-low-attachment 96-well microplates were used (Corning spheroid microplates), uniquely designed with rounded, ultra-low-attachment surfaces to prevent cell adhesion while promoting self-assembly of cells into three-dimensional spheroids as shown in **Figure 2A**. Cell culture medium (DMEM + 10% FBS + 1% penicillin-streptomycin) was replaced daily using a 50/50 approach: replace 50% of the medium to minimize spheroid disturbance. Growth of the spheroids was observed longitudinally via microscopy for as long as they remained viable.

**FIGURE 2:**
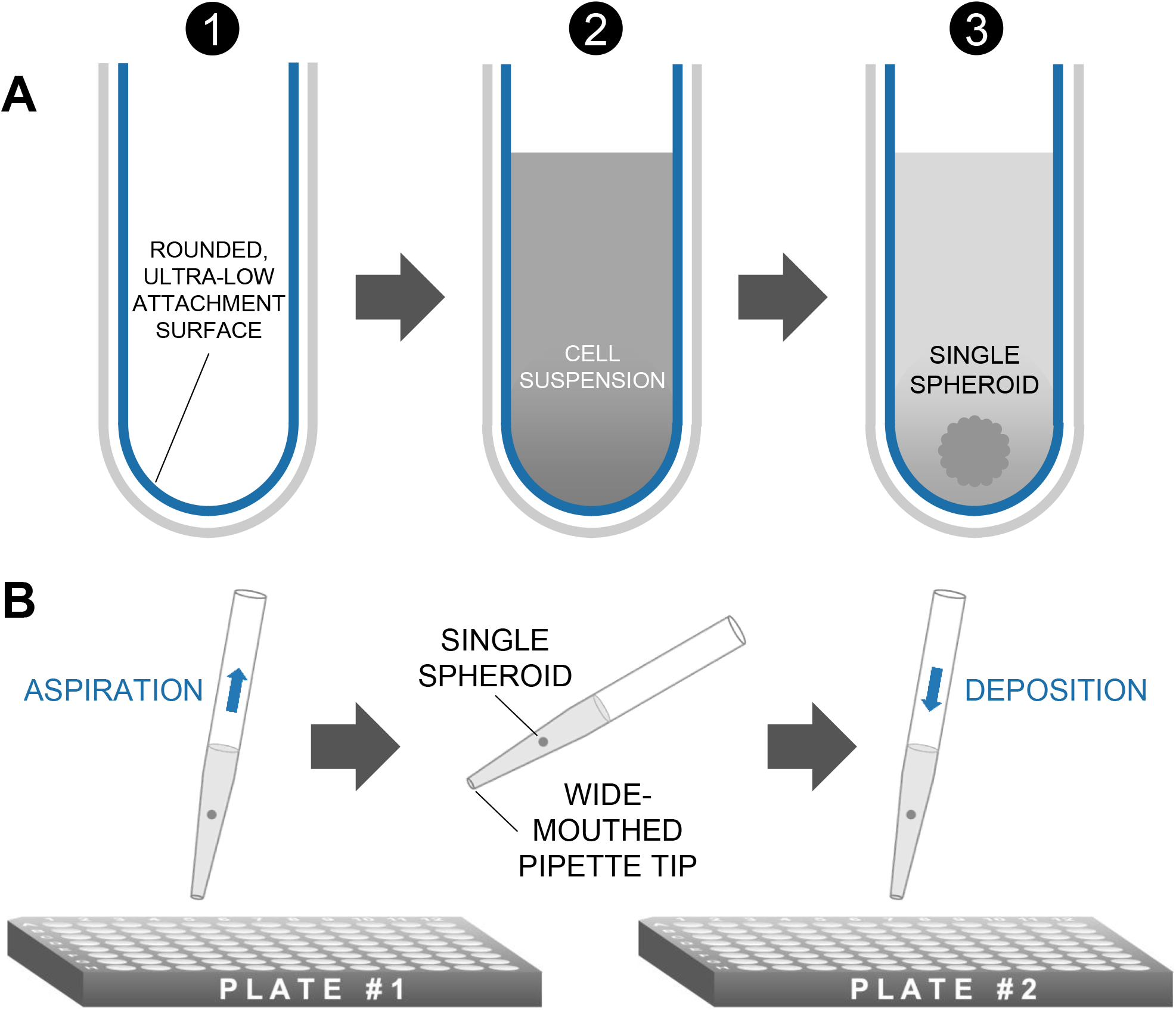
METHOD OF SPHEROID FORMATION. A: Microplates featuring rounded, ultra-low attachment surfaces facilitate the formation of a three-dimensional spheroid of cells from a homogeneous cell suspension. B: Spheroids are transferred in medium via standard micropipettors fitted with wide-mouthed tips.

### CHARACTERIZATION OF SPHEROIDS

Beginning 24 h after seeding, each spheroid was imaged on an inverted Nikon Ti2 microscope, using a 4x objective and an Andor 512×512 EMCCD camera under both brightfield and YFP filter (ChR2-eYFP tag) in 24-h increments, as shown in **Figure 3** and **Suppl. Figures 2-4**. Additionally, 10x and 20x objectives were used to observe the cell structures in more detail and to confirm expression of ChR2-eYFP. The spheroid images shown in **Figure 3** were taken using the 4x objective.

**FIGURE 3:**
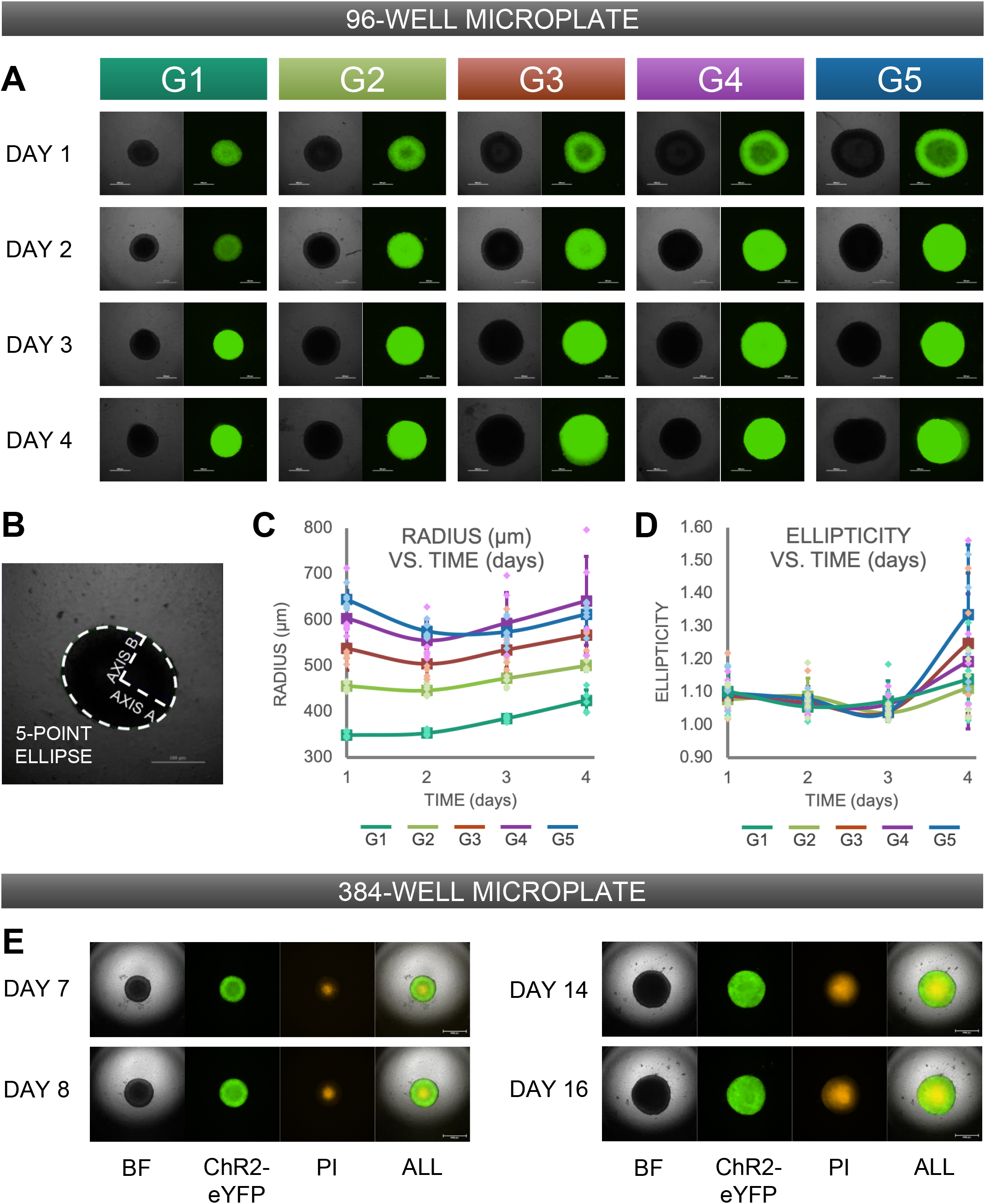
MORPHOLOGICAL CHARACTERIZATION OF 3D SPHEROIDS AS FUNCTION OF CELL DENSITY. A: Depicted are assembled constructs initially seeded at (G1) 2 × 10^4^ cells/well, (G2) 4 × 10^4^ cells/well, (G3) 6 × 10^4^ cells/well, (G4) 8 × 10^4^ cells/well, and (G5) 10 × 10^4^ cells/well. Both brightfield images and fluorescence images (to confirm expression of ChR2-eYFP) were taken every 24 h over a total of 96 h. Scale bar is 0.5mm. B: Standard imaging tools were used to contour a 5-point ellipse to the outline of the imaged spheroid. Axes were then used to calculate average radius and ellipticity for each spheroid over 4 days. C: Radius ((major axis + minor axis)/2) as a function of time. D: Shape (major axis/minor axis) as a function of time. For C and D, the sample size is n = 6 per density and per time point. All data points are overlaid in addition to the mean and standard deviation. E: Sample set of spheroids seeded in 384-well format at 500 cells/well and imaged in brightfield and fluorescence over time. Propidium iodide (PI) was also used to evaluate the viability of the spheroid. Scale bar is 1mm.

Image analysis tools provided by NIS Elements (Nikon microscopes image acquisition software) allowed spatial measurements for major (axis A) and minor (axis B) axes using a five-point ellipse estimation on each spheroid as shown in **Figure 3A**. Size and shape were defined by **Equation 1** and **Equation *2*** respectively, and evaluated every 24 h.

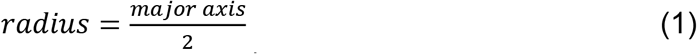

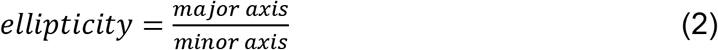

To evaluate spheroid viability, we made use of DNA-binding propidium iodide (Catalog #P3566 Thermo Fisher Scientific). Propidium iodide (PI) is impermeable to cells and thus commonly used to identify dead cells.

After lifting 293Ts and ChR2-293Ts monolayers from culture and resuspending in PI diluted to 2 mg/mL in DMEM, we seeded spheroids at 2 × 10^4^ cells/well. Following imaging every 24 h, medium was replenished using our previously-employed 50:50 method using diluted PI instead of pure DMEM.

Imaging was performed under brightfield as well as appropriate filter cubes to evaluate saturation of PI and ChR2-eYFP expression, respectively. After normalizing the acquisition parameters, we calculated the fraction of PI to total cells for each spheroid. Initially, we ran a simple binary threshold filter on both the PI (numerator) and brightfield (denominator) top-down images for each spheroid. However, given the rounded nature of the wells spheroids are cultured in, we experienced challenges with finding an appropriate threshold to identify the cells in the spheroid (see supplement). Subsequently, for brightfield images, we used an auto-detect ROI feature provided by NIS Elements to identify the general outline of the spheroid and binarize it to the exported image. This revised method for image processing is depicted in **Suppl. Figure 7**. Using this adjusted parameter, we calculated the fraction of PI for each top-down in **Figure 4**. Images of control conditions (no PI stain) for both cell lines are included **in Suppl. Figure 6**.

**FIGURE 4:**
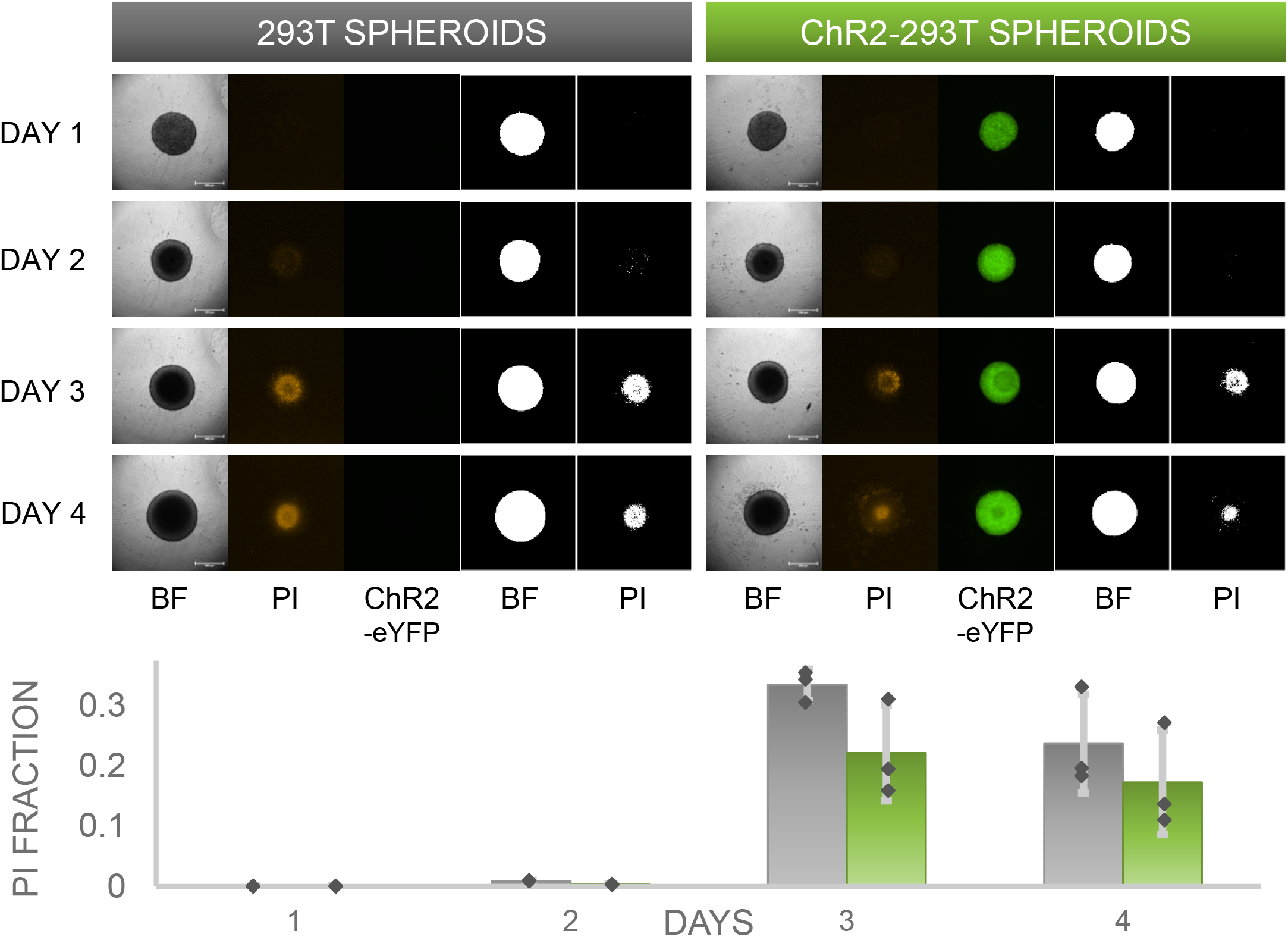
SPHEROID VIABILITY. To evaluate the viability of spheroids in culture over time, spheroids at seeding density 2 × 10^4^ cells/well were cultured in a sterile propidium iodide (PI) solution of concentration 2 mg/mL in DMEM. Spheroids were imaged every 24 h under brightfield and fluorescence to reveal the PI uptake localization in the necrotic core of the spheroid. In both 293T spheroids and ChR2-293T spheroids top-down images, we observed an increase in the fraction of PI pixels to total spheroid pixels (as determined from brightfield) after day 2. These fractions were quantified over time, n = 3 spheroids per time point per group, all data points shown.

### CULTURE OF STEM-CELL-DERIVED CARDIOMYOCYTES

Human induced pluripotent stem-cell-derived cardiomyocytes, hiPSC-CMs (Catalog #R1017, Fujifilm - Cellular Dynamics) were thawed and plated according to the supplied protocol. Prior to thawing, a 50 μg/mL solution of fibronectin diluted in PBS was applied to all wells / plates which were incubated overnight at 37°C. Maintenance medium was replaced every 48 h. For initial and integration experiments, cells were plated in 96-well glass bottom plates at 50 × 10^3^ cells/well. For macroscopic experiments, hiPSC-CMs were seeded into 14 mm glass-bottom dishes at a seeding density of 270 × 10^3^ cells/well.

### DEPOSITION OF SPHEROIDS ONTO CARDIAC SYNCYTIA

ChR2-HEK spheroids at 2 × 10^4^ cells/well initial seeding density and cultured for 24 hours were transferred via pipetting onto hiPS-CM syncytia, plated 5 days before, or 7 days prior if transfection was applied. The fragile nature of the spheroids called for careful attention during handling, such as a wide-mouthed pipette tip as well as a slow aspiration and deposition technique when a spheroid was transferred from one well to another, as shown in **Figure 2B**.

### IMMUNOCYTOCHEMISTRY

Immunocytochemistry was used to visualize the spheroid-syncytia constructs. The ChR2-expressing spheroids have an eYFP fluorescent reporter. For the hiPSC-CMs, we used the monoclonal anti-α-actinin antibody (Catalog #A7811, Millipore Sigma) to label cardiomyocyte sarcomeres and Hoechst (Catalog # H3570, Thermo Fisher Scientific) to label the nuclei. Following rinsing with PBS, cells were permeabilized in 0.2 Triton X-100 in 5% FBS and two-stage antibody labeling was applied to image the samples on an inverted microscope.

### FUNCTIONAL EXPERIMENTS USING ALL-OPTICAL ELECTROPHYSIOLOGY

Functional experiments were conducted using all-optical electrophysiology [17; 24]. Briefly, all functional experiments were performed in Tyrode’s solution: (in mM): NaCl, 135; MgCl_2_, 1; KCl, 5.4; CaCl_2_, 1.8; NaH_2_PO_4_, 0.33; glucose, 5.1; and HEPES, 5 adjusted to pH 7.4. For voltage measurements, 24h grown ChR2-HEK spheroids of 2 × 10^4^ cells per well were deposited onto hiPSC-CM syncytia in a 96-well format. After 12 h of integration, the samples were labeled with small-molecule near-infrared voltage-sensitive BeRST1 dye, courtesy of Evan W. Miller [25] at 1 mM. For measurements of intracellular calcium, we used either Rhod-4AM ((AAT Bioquest, Sunnyvale, CA) at 10 mM of the genetically encoded calcium indicator (GECI) R-GECO, expressed in the hiPSC-CMs before the deposition of the spheroids. R-GECO expression was done with lipofectamine 3000 using the Addgene plasmid CMV-R-GECO-1.2 at 400 ng per sample (Catalog #45494, Addgene), courtesy of Robert Campbell [26]. All microscopic measurements were done on an inverted microscope Nikon Eclipse Ti2 at 20x. Blue light 470 nm via a digital micromirror device DMD (Polygon 4000, Mightex, Toronto, Canada) were used to stimulate the spheroids, 20ms pulses at <1mW/mm^2^ and using 0.5 Hz-2.0 Hz in order to trigger excitation in the iPSC-CMs. Excitation for voltage imaging with BeRST1 was at 660 nm and emitted light was collected using a long-pass filter at 700 nm, excitation for calcium imaging with Rhod-4AM and R-GECO was at 535 nm and emission was collected at 605 nm, in both cases with an iXon Ultra 897 EMCCD camera (Andor Technology Ltd., Belfast, UK), run at 200 fps. To avoid potential spectral overlap in the R-GECO measurements, we used patterned light, where the DMD stimulated region and the recorded region were physically different.

### PROBING SPHEROID / SYNCYTIA COUPLING

After deposition of ChR2-HEK spheroids (2 × 10^4^ cells/well, 24 hours) onto R-GECO-transfected hiPSC-CMs (10 days in culture including transfection), samples were transferred to an on-stage microscope incubator and probed for responsiveness to blue light (20 ms at 470 nm <1mW/mm^2^) pulsed at 0.5 Hz-2.0 Hz. Use of the incubator ensured temperature control at 37°C throughout the length of the 10-12 h experiment.

Functional tests with R-GECO as described in detail before were performed and repeated in two-hour increments over 10 hours for a total of 6 time-probes. This procedure allowed us to determine the emergence of system responsiveness to optical stimulation by examining the number successfully stimulated samples over time.

### MACROSCOPIC OPTICAL MAPPING

For imaging the macroscopic waves triggered by the spark-cell spheroids, we deposited the ChR2-293T spheroids (2 × 10^4^ cells/well, 24 h) onto hiPSC-CM samples plated in 14 mm glass-bottom dishes after 24 h of integration.

The widefield optical mapping of cardiac excitation wave was built around an MVX10 MacroView, Olympus, Japan system. We used a high-speed CMOS camera (Basler, Ahrensburg, Germany) to track wave dynamics. Either Rhod-4AM or R-GECO were used for these measurements. Continuous fluorescence excitation light with an irradiance of 0.28 mW/mm^2^ was generated by a collimated green LED (M530L3, Thorlabs, USA), which obliquely illuminated the hips-CMs dish from underneath. Pulsed optical pacing light with an irradiance of 0.35 mW/mm^2^ was generated by a weakly focused LED (M470L4, Thorlabs, USA), which was driven at 0.5 Hz (or 1Hz) with a pulse duration of 10 ms. Microscopic and macroscopic recordings were processed using Matlab software for filtering and visualization [18; 27].

### GENE EXPRESSION ANALYSIS BY qPCR

Cells were plated on 96-well format. Two days post plating, cells were harvested for RNA extraction and mRNA levels were detected and quantified using Power SYBR^TM^ Green Cells-to C_T_^TM^ Kit (Cat. 4402953, Invitrogen) according to the manufacturer’s protocol. qPCR analysis was performed on a Quantstudio 3 Real-Time PCR System (ThermoFisher Scientific) with the QuantStudio Design and Analysis Software (ThermoFisher Scientific). Cx43 (Fw_GGTGGTACTCAACAGCCTTATT; Rev_ACCAACATGCACCTCTCTTATC) gene expression quantification was normalized to expression of housekeeping gene GAPDH (Fw_GGAGCGAGATCCCTCCAAAAT; Rev_GGCTGTTGTCATACTTCTCATGG) using standard ΔΔCt method.

### PROTEIN QUANTIFICATION BY WES

Cells were lysed for total protein in 96-well format using the Qproteome Mammalian Protein Prep Kit (Cat. 37901, Qiagen). Protein lysates were loaded onto and analyzed using the Wes^TM^ (ProteinSimple) capillary-based system for protein quantification. Proteins of interest were probed using antibodies specific to Cx43 (ab11370, Abcam) and GAPDH (ab181602, Abcam). Changes of Cx43 protein levels were presented as normalized to GAPDH.

## Results

### MANUFACTURING OF “SPARK-CELL” SPHEROIDS

Our goal was to scale the TCU strategy up, **Figure 1A**, into an easy-to-handle, configurable three-dimensional structures of ChR2-expressing cells for use in high-throughput drug screening of cardiac tissue. Such “spark-cell” spheroids composed of optogenetically-transformed live cells can be positioned atop hiPSC-CM syncytia, as shown in **Figure 1B,** to mediate optical pacing.

While various methods exist to form three-dimensional cell constructs, including 3D bioprinting with hydrogels and photopolymerizable materials [28; 29], we opted for spheroid formation via self-aggregation, due to its simplicity. Indeed, it has been demonstrated that cells deposited in agarose molds can self-assemble into three-dimensional tissue spheroids [30; 31]. We fabricated our ChR2-HEK spheroids in commercially available 96-well microplates with ultra-low attachment walls, as in **Figure 2A**. Rounded wells treated with adhesion deterent force cells to adhere to one another, forming a spheroid over time. This assembly is further facilitated by gravity. It was possible to use wide-mouth pipette tips to handle these spheroids, i.e. to carefully lift, transfer, and to deposit them on top of hiPSC-CM monolayers (gown in 96-well plates), as shown in **Figure 2B**.

Optimization of seeding cell density included considerations for easy handling (the spheroids had to be large enough) and avoiding the formation of a significant necrotic core (not too large). Spheroids with initial seeding density between 10^4^-10^5^ cells/well yielded viable and functional spheroids recognizable by the human eye.

### CHRARACTERIZATION OF SPHEROIDS

Different seeding densities from 2 × 10^4^ cells / well to 10^5^ cells / well in increments of 2 × 10^4^ cells / well were explored. To register how spheroids evolved structurally over time, brightfield images were captured every 24 h, along with images of YFP to monitor the expression of ChR2-eYFP, **Figure 3A**. Over the first days, the spheroids maintained uniform shape and grew in size consistent with the proliferation property of HEK cells. Beyond 96 h in culture, most of these spheroids were prone to developing aberrant growths and ultimately became more fragile.

For morphological analysis of the spheroids, standard image analysis was used and ellipses from five points were formed using 4x images, as illustrated in **Figure 3B**. From each 5-point ellipse, major and minor axes were calculated to yield measurable parameters, such as radius and ellipticity. Data is depicted for both size and shape as a function of time for individual density groups (n=6 per density and per day) in **Figures 3C-D**. Spheroids at all densities became more compact over the first 48 h, then grew larger due to proliferation. Furthermore, spheroids at all densities stayed close to a perfect cicular shape for the first 3 days, however after 72 h in culture their elipticity increased. A two-way ANOVA showed significant differences in the size of spheroids over time as well as the shape of spheroids over time (p < 0.01).

We observed that to control better shape and size, it was important to scale down the initial seeding density. To do so, we utilized a smaller well format (384-well ultra-low-adhesion microplates) in conjunction with lower cell densities from 0.5-8 × 10^3^ cells/well. Shown in **Figure 3E** are a sample set of spheroids that were initially seeded in 384-well ultra-low-adhesion plates at a density of 500 cells/well. These smaller spheroids did not demonstrate any stages of compression, but rather steadily grew in size throughout time in culture. Importantly, these spheroids at 40-fold sparser initial density in fact maintained their round shape into day 7 of culture. The sample shown in **Figure 3E** was able to sustain a consistent shape over 16 days of culture.

To track spheroid viability over time, they were formed resuspended in DMEM, containing PI at 2 mg/mL, as Zhao et al. have shown the safe long-term use of PI [32]. Every 24 h, cells were imaged and medium replaced with the same PI dilution to be sure concentration gradients of the PI were maintained. This characterization experiment generated the images for **Figure 4** in which WT 293T and ChR2-infected 293T spheroids were seeded at 2 × 10^4^ cells/well monitored over time. Binarization of the PI images revealed a quantitative PI fraction for each cell line and day (n = 3 for each condition), summarized in **Figure 4.** Both the smaller and the larger spheroids developed a necrotic core with time in culture, as per **Figure 3E** and **Figure 4**, surrounded by viable cells, thus not affecting their acute use to pace cardiomyocytes.

### GAP JUNCTIONS AND THE “SPARK-CELL” SPHEROIDS

Immunocytochemistry and confocal imaging helped visualize the coupling of the spheroids with cardiomyocyte layers (**Figure 5A**). Of interest was the coupling mechanism between the ChR2-spheroid and hiPSC-CM syncytia in the this larger-scale TCU approach. We expected that gap junctional proteins, such as Cx43 (abundantly expressed in ventricular cardiomyocytes), play a role, even though the HEK cells are known to have minimal amounts of Cx43. We used small-sample qPCR and protein quantification assays to probe for GJA1/Cx43 expression levels, normalized by GAPDH transcript/protein, respectively. As expected, the lysed monolayers and spheroids of WT-HEK and ChR2-HEK cells had an order of magnitude lower GJA1 mRNA levels (normalized to GAPDH) compared to the human iPSC-CMs, **Figure 5B.** Yet, these were detectable, and in the spheroids, the ChR2 expression seemed to increase GJA1 mRNA. At the protein level, ChR2-HEK also showed a trend towards increased normalized Cx43 protein levels compared to the WT HEK, as it has been seen earlier [23]. The presence of Cx43 in the “spark-cell” spheroids suggests gap junctional contribution to the their integration with the human iPSC-CMs.

**FIGURE 5:**
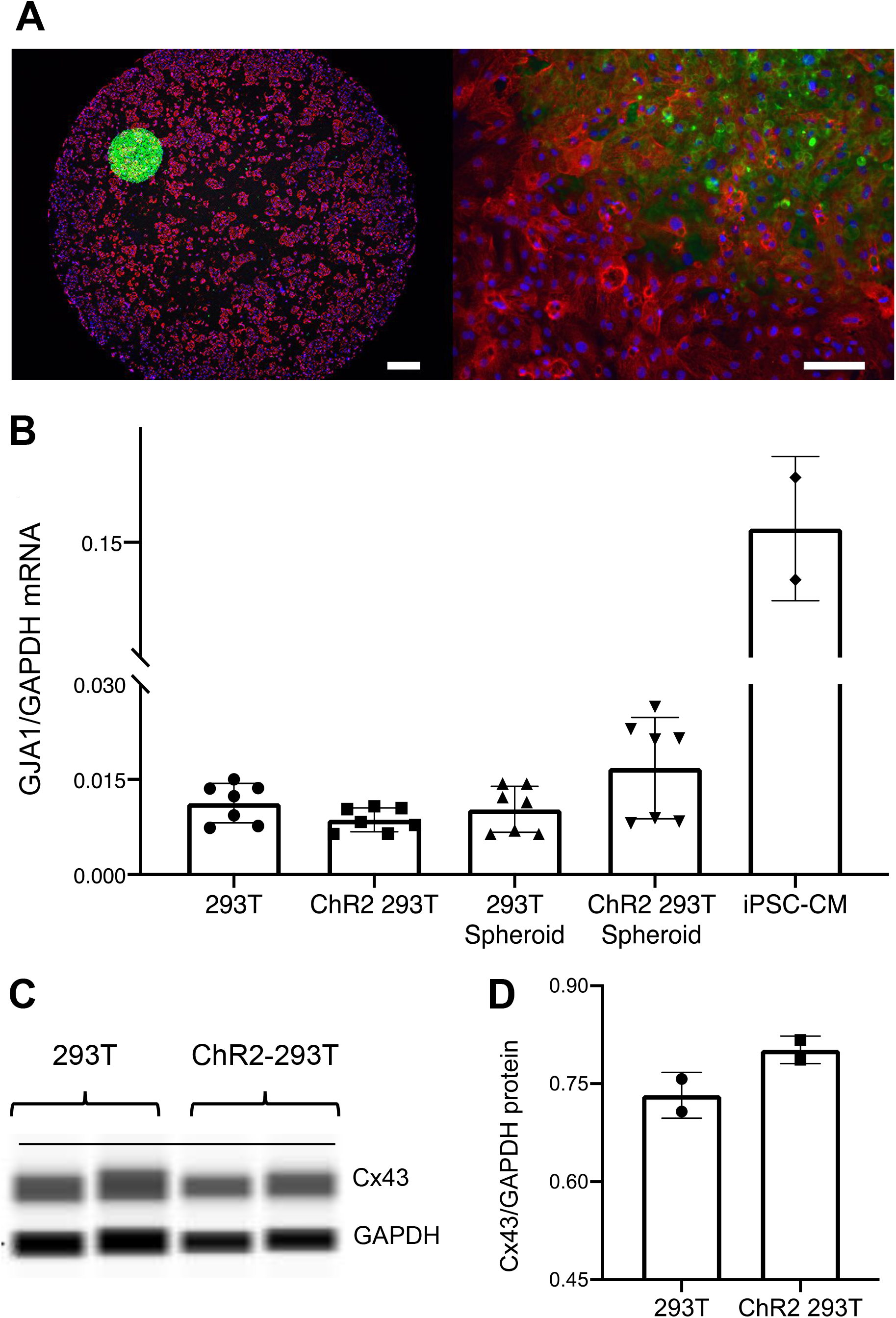
SPHEROID DEPLOYMENT AND COUPLING TO HUMAN IPS-CM SYNCYTIA. A: For acute functional studies, spheroids were deposited onto pre-plated hiPSC-CM syncytia. A: To investigate system structure, samples were labeled with fluorescent dyes. Alpha-actinin (red) identifies the actin filaments in the sarcomeres of the myocytes; Hoechst (blue) labels the cell nuclei; eYFP (green) reporter identifies the expression of ChR2 which is localized in the HEK cell membrane. Left scalebar is set to 0.5 mm. Right scalebar is at higher magnification set to 0.1 mm. B: Quantification of gap junction (GJA1) gene expression via qPCR (normalized by GAPDH). Shown are data from n = 7 independent samples of WT HEK and ChR2-HEK cells grown in monolayers and in spheroids, as well as n = 2 samples of hiPSC-CMs. C and D: Western blot (by electrophoresis-based Wes) quantification of Cx43 gap junctional protein from WT HEK and ChR2-HEK cells, normalized by GAPDH protein, n = 2 samples per case.

### DEMONSTRATION OF FUNCTIONALITY

To optimize the stability of the spheroids, we determined that the most impactful constraints were early time in culture and low initial seeding density, e.g. 2 × 10^4^ cells / well. Hence, all functional testing experiments were performed at this cell density after 24 h upon depositing the spheroids onto 96-well monolayers of hiPSC-CMs (pre-cultured for 5 days post-thaw) as described in **Figure 2B**.

Typical all-optical electrophysiology experiments (as in **Figure 6**) were done 12 h post spheroid deposition onto hiPSC-CMs grown in 96-well plates. For optical confirmation of pacing, we used genetically-encoded calcium sensor (GECI) R-GECO, expressed in the iPSC-CMs, or near-infrared voltage-sensitive dye, BeRST1, both of which are spectrally-compatible with the ChR2 actuator [13; 17]. To further suppress potential cross-talk when using calcium recordings with R-GECO, we spatially patterned the light to confine the excitation near the spheroid and optically recorded from a nearby region, **Figure 6B**. Such restrictions were not necessary when using the red-shifted voltage-sensitive dye.

**FIGURE 6:**
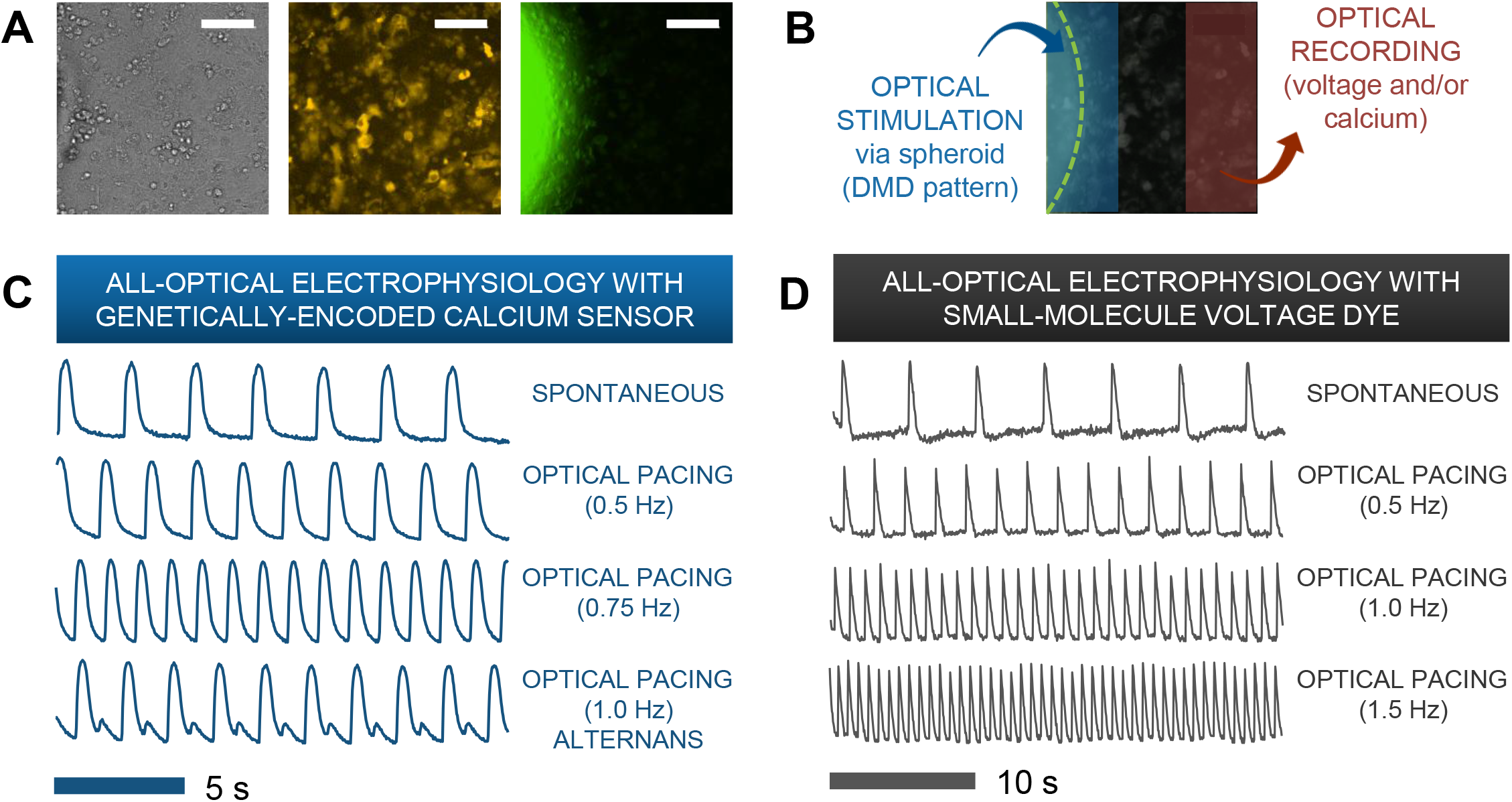
FUNCTIONAL RESPONSES FROM HUMAN IPS-CMS UPON SPARK-CELL CLUSTER PACING OBTAINED BY ALL-OPTICAL ELECTROPHYSIOLOGY. A: A typical field-of-view for microscopic optical stimulation experiments featuring brightfield and corresponding fluorescence images. The middle image was produced using an mCherry filter and demonstrates expression of optogenetic sensor for calcium (R-GECO) in the iPSC-CM monolayer. The rightmost image was produced with a YFP filter and shows a portion of the ChR2-eYFP spheroid. Scale bar is 0.1mm. B: For recordings, the field-of-view was separated into two non-overlapping areas - patterned optical stimulation including the spheroid and optical recording including only iPSC-CMs labeled with voltage or calcium indicators. C: Calcium transients from R-GECO-expressing iPSC-CMs showing spontaneous activity and optical pacing (via the spheroid) at three different frequencies after 7 h of spheroid introduction. While a 1:1 stimulation-to-response is observed for optical pacing at 0.5 Hz and 0.75 Hz, overdrive pacing at 1.0 Hz results in failure of iPS-CMs to fully restore calcium stores between stimulation events, hence calcium alternans are seen. D: Optical pacing can also be confirmed via spectrally-compatible small-molecule voltage dye. This sample shows a 1:1 relationship between stimulation and response for pacing frequencies up to 1.5 Hz after 12 h of spheroid integration.

**Figures 6C-D** show traces of intracellular calcium (via R-GECO) and action potentials (via BeRST) from spontaneous and paced activity via “spark-cell” spheroids. Successful pacing was confirmed when samples followed 1:1 the selected frequencies over at least 10 consequitive beats. Incomplete capture sometimes resulted in alternans or 2:2 response [27], as shown in **Figure 6C**.

To confirm that the “spark-cell” spheroids are capable of pacing cells in larger cm-scale cardiac samples and to map the waves triggered by the spheroids, we also performed some macroscopic optical mapping experiments in 14 mm dishes, **Figure 7**. For these experiments, the spheroids were positioned on top of the cardiomyocytes 24 h before the imaging. Indeed, we were able to register spheroid-triggered excitation waves using either R-GECO or Rhod4-AM calcium sensor, as shown in **Figure 7**. Light was aimed at the spehroids but not focused on them. When more than one spheroid was used, as in **Figure 7A**, a dominant pacemaker emerged, driving the cardiac syncytium on each beat.

**FIGURE 7:**
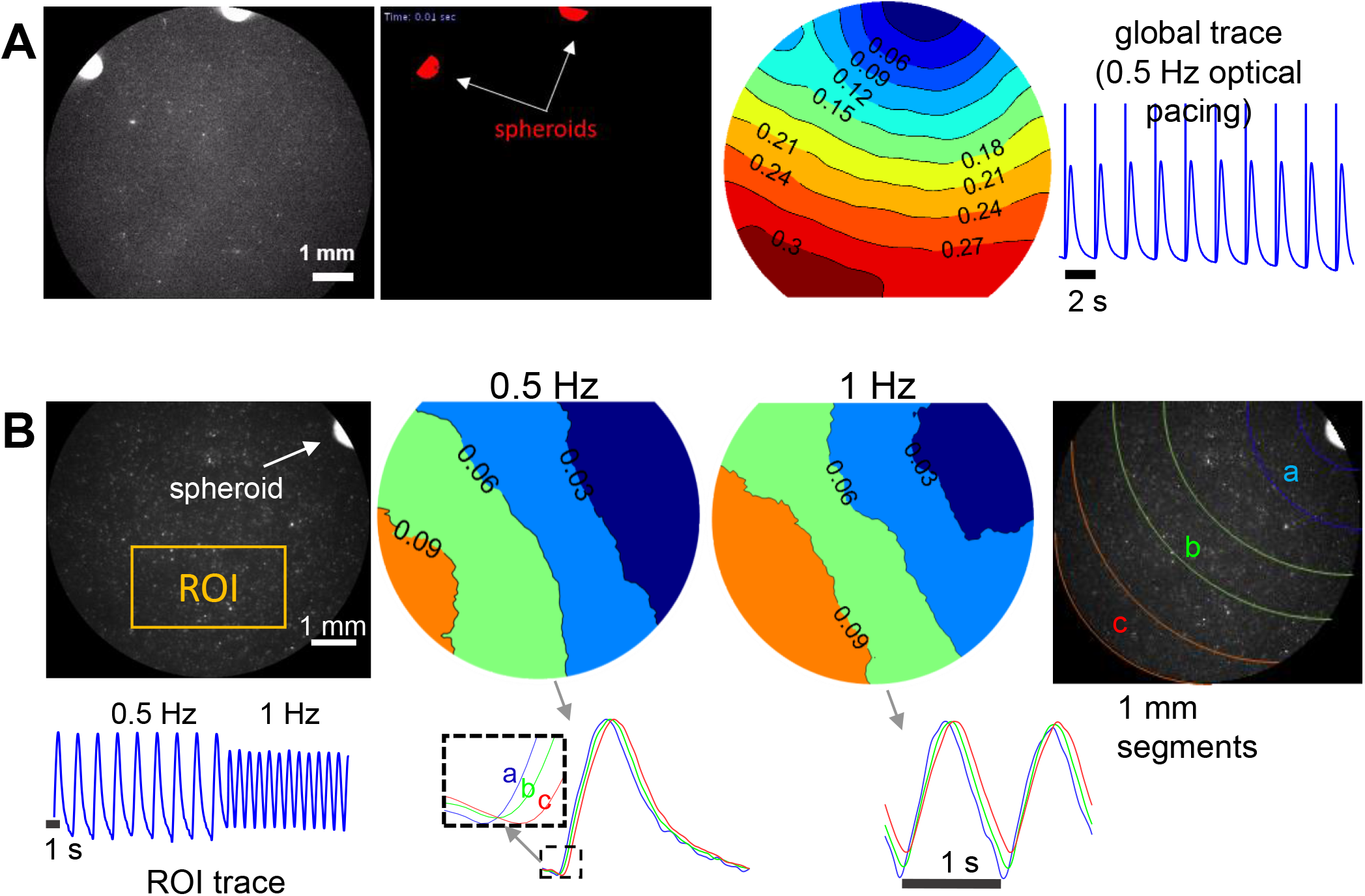
MACROSCOPIC OPTICAL MAPPING OF OPTICALLY-TRIGGERED SPHEROID-MEDIATED CALCIUM WAVES IN HUMAN IPS-CMS. A: Calcium waves (by Rhod-4) triggered by optical light pulses at 0.5 Hz through a pair of spheroids after 24 h integration. Activation map shows the origin of early activation (blue) and isochrones are 0.03 s apart. Global calcium transients and the superimposed optical pulses are shown on the right. B: Calcium waves by optogenetic sensor R-GECO, triggered at 0.5 Hz and at 1 Hz through a single spheroid (after 24 h integration). In the activation maps (isochrones 0.03 s apart) slight wave-slowing is visible at the higher pacing frequency. At the bottom, an ROI trace is shown for the uninterrupted pacing and traces from different segments away from the spheroid illustrate sequential activation (a, b, c). Scale bar is 1 mm.

### TIMING OF INTEGRATION

After confirming spheroid functionality, we sought to probe the earliest emergence of TCU-based coupling in our system. To do so required the approximate timeline of experimental preparation shown in **Figure 8A,** followed by testing a set of 18 x iPSC-CM samples with ChR2-HEK spheroids over 10 hours as shown in **Figure 8B**. Emergence of pacing was registered in some samples as early as 2 h after first depositing the spheroids onto the hiPS-CM syncytia. Integration between spheroid and syncytia increased over the experimental timeline, with 6 samples out of 18 showing successful pacing via spheroid at the end of 10 h. Over multiple cultures, we estimated the percentage of samples that followed optical pacing at 6 h and at 24 h after introducing the spheroids. As shown in the inset of **Figure 8B**, samples from three separate iPSC-CM cultures (n = 16, 18, and 20 for total n = 54) yielded successful pacing at 24% after 6 h and 81% after 24 h. As done in other studies [15; 33], we considered that spheroid positioning via magnetic nanoparticles may even further speed up integration. As described in the *Supplementary Information*, in this study, this was no successful, yet this approach may offer benefits with further optimization. Overall, we concluded that the “spark cell” spheroid approach has the potential to yield functional optical pacing significantly earlier than the standard direct viral or liposomal transductions of the cardiomyocytes, which typically require 48 h.

**FIGURE 8:**
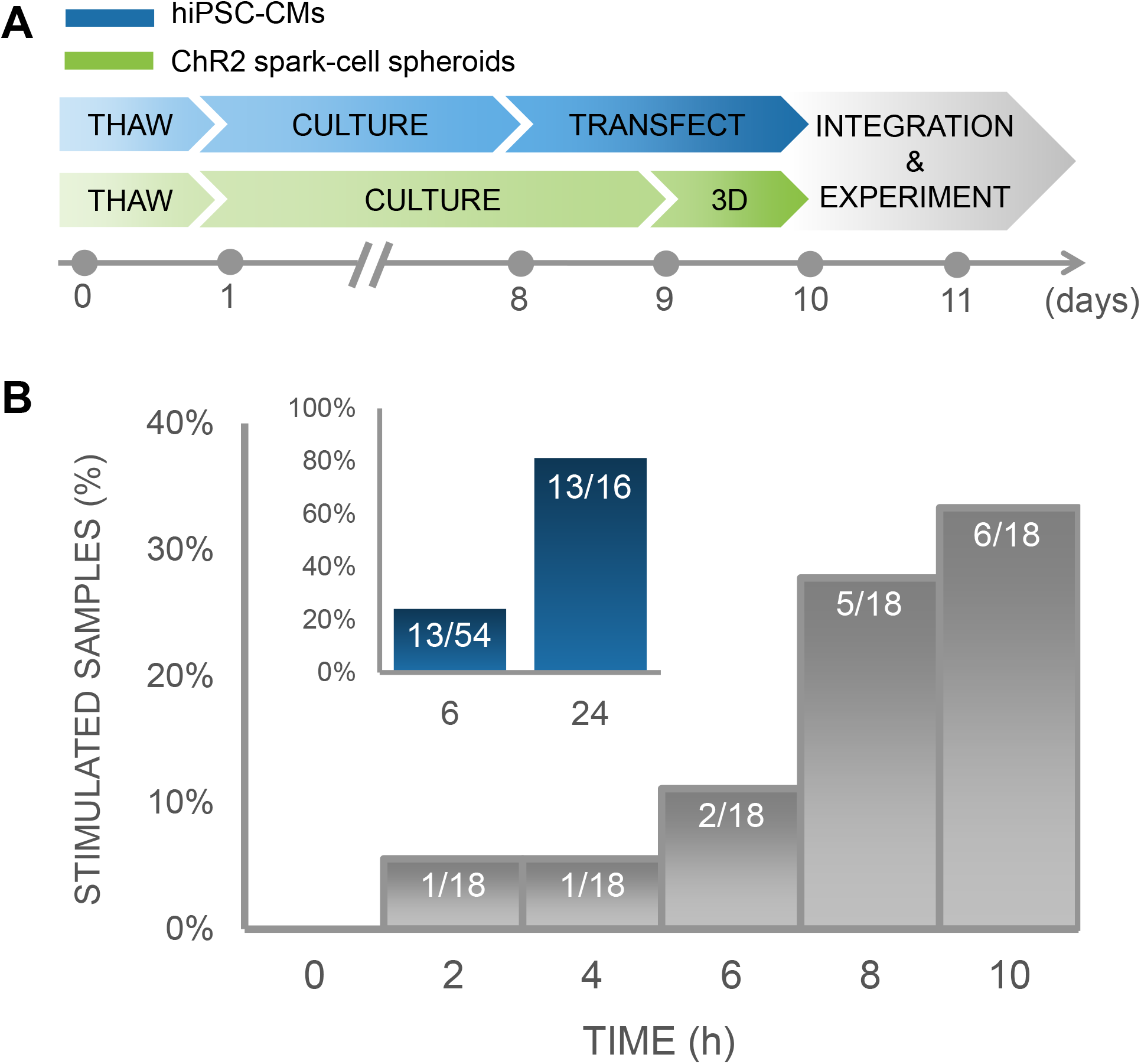
TIME COURSE OF INTEGRATION OF SPARK-CELL CLUSTERS WITH HUMAN IPS-CM MONOLAYERS FOR PACING CAPTURE. A: Experimental preparation timeline including separate treatment of cell lines (hiPSC-CM in blue and ChR2 293Ts in green) before acute functional studies. B: For an early integration study, 18 samples were tested and the percentage of those samples from which we observed full pacing capture was plotted every 2 h. We observed an increase in the number of optical pacing responders over 10 h. The inset (blue), represents the combined results over multiple cell cultures, summarized as a percent of samples that responded to optical pacing at 6 h (n = 54) and at 24 (n=16) after spheroids were seeded onto hiPSC-CM syncytia.

## DISCUSSION AND CONCLUSIONS

High-throughput drug screening, and specifically cardiotoxicity testing, can benefit from optical actuation and sensing methods that offer contactless and scalable interrogation of cardiomyocytes. Optogenetic approaches play a key role in the development of high-throughput all-optical electrophysiology. Some of the limitations associated with (opto)genetic modification of cells include potential interference with their innate functional responses and the time needed for the genetic modification to produce the protein of interest.

In this study, we sought to biomanufacture “spark-cell” spheroids, usable as a “reagent” that eventually can be stored, transported and deployed on site to confer optical pacing of cardiac tissue. Such “spark-cell” spheroids are amenable to automation and can be handled robotically. When deposited onto human iPSC-CM syncytia, they create a spatially-localized pacing site. Our results show that this approach may offer faster integration for acute pacing experiments (as early as 2 h post-deposition in some cases) compared to direct genetic modification of the cardiomyocytes. The mechanism of optical pacing is based on the TCU approach and likely involves the formation of close contact and gap-junction mediated ion currents between the spheroid and the responding cardiomyocyte syncytia.

Additional optimizations can speed up the contact formation even further using magnetic assemblies, microfluidic deposition or other augmentations. In the *Supplementary Information*, we describe several approaches that we tried and they yielded negative results, including attempts to control proliferation of the cell spheroids (as an alternative to simply reducing size), magnetically-assited positioning and proof-of-principle cryopreservation and thawing. Including such negative results may be useful to inform others what we have tried and save them time but also because with further optimization, these ideas may lead to significant improvements of the approach. Drawing on published work on spheroids made out of cardiac progenetor cells, cardiac fibroblasts or cardiac myocytes [34; 35; 36; 37], the proposed optogenetic approach can facilitate the asembly of modular designer tissues with space-patterned pacemakers in 3D. It can find translational applications in the pharmaceutical industry for acute and chronic drug testing. The methodology can also be useful for aiding the long-term maturation of human iPSC-CMs via optical pacing for regenerative purposes.

## Acknowledgements

This work was supported in part by an NIH grant R01HL144157 and grants from the National Science Foundation EFMA 1830941 and PFI 1827535 to E.E. as well as a SEAS SUPER Fellowship and Clare Boothe Luce Research Fellowship awarded to C.C.

